# A new microbiological weapon against lepidopteran pests

**DOI:** 10.1101/2022.05.06.490868

**Authors:** Vyacheslav V. Martemyanov, Yuriy B. Akhanaev, Irina A. Belousova, Sergey V. Pavlusin, Maria E. Yakimova, Daria D. Kharlamova, Alexander A. Ageev, Anna N. Golovina, Sergey A. Astapenko, Alexey V. Kolosov, Grigory G. Ananko, Oleg S. Taranov, Alexander N. Shvalov, Sergey A. Bodnev, Nikita I. Ershov, Inna V. Grushevaya, Maxim A. Tymofeev, Yuri S. Tokarev

## Abstract

Nowadays researchers provide more and more evidence that it is necessary to develop an ecologically friendly approach to pest control. This is reflected in a sharp increase in the value of the biological insecticide market in recent decades. In our study, we found a virus strain belonging to the genus *Cypovirus* (Reoviridae); the strain was isolated from *Dendrolimus sibiricus:* that possesses attractive features as a candidate for mass production of biological agents for lepidopteran-pest control. We describe morphological, molecular, and ecological features of the new *Cypovirus* strain. This strain was found to be highly virulent to *D. sibiricus* (half-lethal dose is 68 occlusion bodies per second-instar larva) and to have a relatively wide host range (infects representatives of five families of Lepidoptera: Erebidae, Sphingidae, Pieridae, Noctuidae, and Lasiocampidae). The virus strain showed a strong interaction with a nontoxic adjuvant (optical brightener), which decreased the lethal dose for both main and alternative hosts, decreased lethal time, and may expand the host range. Moreover, we demonstrated that the insecticidal features were preserved after passaging through the most economically suitable host. By providing strong arguments for possible usefulness of this strain in pest control, we call on virologists, pest control specialists and molecular biologists to give more attention to the *Cypovirus* genus, which may lead to new insights in the field of pest control research and may provide significant advantages to compare with baculoviruses and Bacillus thuringiensis products which are nowadays main source of bioinsecticides.

**Significance statement:** Within this article we are describing unique set of features of newly discovered cypovirus strain which possess by significant premises for modern biological insecticides requirements: high potency, universality, true regulating effect, flexible production (possibility to choose host species for production), interaction with enhancing adjuvants, ecologically friendly. Basing on genome alignment we suggest that increasing of host range of new strain is the sequence of evolutionary event which was occurred after coinfection of different CPV species within same host. This finding open new perspective to consider CPVs as perspective agent of biocontrol products.

## Main

The large increases in the scale of agriculture and forestry in the last century—which are necessary for the functioning of human populations at their current density—stimulate the emergence of new pest species. Thus, humans will be strongly dependent on pest management if they want to keep their population density while maintaining their standard of living. The simplest way to control the population size of pests is the use of chemical insecticides. Major shortcomings of these insecticides are as follows: the duration of the effect is inadequate (i.e., this is not true regulation), and chemical insecticides are not environmentally friendly. Rapid development of pests’ resistance to chemical insecticides is another major disadvantage of chemical control and forces researchers to keep designing new insecticides frequently. In the meantime, an alternative way to manage the density of pests is the use of biological agents, such as natural enemies or diseases. The latter approach is gaining popularity in this century. The attractiveness of this method is based on recent studies showing the importance of clean environment (including good-quality food) for human health, and in this regard, biologicals are usually environmentally friendly. This fact strongly contributes to new developments in the production of so-called organic foods (i.e. those produced without chemical pesticides), which are becoming increasingly popular in many countries worldwide. Nonetheless, the direct effect of organic foods on human health is still debated^1^.

Despite discussions about the association between human health and organic-food consumption, we clearly see a substantial rise of the bioinsecticide market: from modest size in the 1990s to a multibillion-dollar business today that is growing at a much faster rate than the conventional crop protection industry (https://www.bpia.org/markets-for-biological-products-global-market-landscape/). The market size of agricultural biologicals was estimated to be 8.8 billion US dollars in 2019 and is projected to grow at a CAGR of 13.6% and reach 18.9 billion US dollars by 2025 (Agricultural Biologicals Market report). Detailed analysis of the structure of this market reveals unequivocal dominance of *Bacillus thuringiensis* (Bt) on the basis of the offered products^2^. The key reasons for this state of affairs are the possibility of cultivation of this bacterium on a commercial medium and the broad specificity of bioinsecticides, which allows users to apply the same bioinsecticide against different representatives of one order (e.g., against Lepidoptera, Diptera, or Coleoptera). The serious drawback of Bt-based products is the absence of a true pest-population-regulating effect that reduces application frequency of such bioinsecticides against polyvoltine pest species in season. Viral bioinsecticides based on entomopathogenic viruses (mostly representatives of genera *Nucleopolyhedrovirus* [NPV] and *Granulovirus*; Baculoviridae) are free from this disadvantage owing to vertical transmission via survival of the hosts after infection^3^; in combination with extremely high specificity to a target host, this feature has allowed these products to secure a good percentage of the market^2^. The basic limitations of virus-based products are the current impossibility of mass cultivation in cultured cells (because of two-step ontogenesis of such viruses and high prices of cell culture media^2^) and very high specificity. This situation necessitates breeding a specific host species for each viral product (i.e., a lack of universality of a given product in a shifting market).

In this article, we describe a new strain of an RNA virus from the *Cypovirus* (CPV) genus (Reoviridae family), which has been shown to possess important features in terms of its effectivity, productivity, host range, production prospects, interaction with an adjuvant, and overall good potential for the organic-food industry. CPVs, also known as cytoplasmic polyhedrosis viruses, usually cause chronic infection in the gut of insects^4, 5^ and often participate in a coinfection with baculoviruses^6, 7^. These features attract much less interest of researchers as compared with typical “killers” in the world of pathogens such as the *Baculoviridae* family (16000 articles for search term “Baculovirus” vs. 200 articles for search term “Cypovirus”). Such intense research attention has given rise to a new scientific field related to baculoviruses: Baculoviridae-based in vitro expression systems, which enable investigators to accumulate target proteins in an applicable amount^8^. The balance of research attention to these two groups of occlusion body–forming viruses is dramatically skewed toward the Baculoviridae side if we compare the pest management products in terms of these viruses: many nucleopolyhedrovirus-based and granulovirus-based bioinsecticides^9, 10^ vs. a few bioinsecticides based on *Dendrolimus punctatus* CPV^10^.

The main aim of the present study was to describe morphological, molecular, and biological features of a new strain of CPV isolated from *Dendrolimus sibiricus* (Lepidoptera, Lasiocampidae), which is a major forest pest in boreal forests of Asia. The isolate was designated as DsCPV. Another aim was to study the host range of this virus strain including a host species that is already adapted to artificial inexpensive mass cultivation. Finally, we were interested in assessing possible synergy of the interaction of this virus with a nontoxic adjuvant, such as an optical brightener, which usually has strong synergistic effects with NPV representatives on insect mortality^11^. These data will facilitate the development of a relevant virus product for field application.

### 1. Virus isolation

The virus was isolated from dead *D. sibiricus* larvae collected in 2020 in fir-cedar forests located at the foothills of the Eastern Sayan (55.059652°N, 96.050832°E). The first identification of the virus was performed by microscopic analysis of a rough imprint of the dead larvae contents on a histological slide. For subsequent manipulations with the virus, the dead larvae were crushed in distilled water, and then polyhedral occlusion bodies were separated from rough debris by filtration through gauze, with subsequent centrifugation at 20000 ×*g* for 20 min. The obtained virus samples were then used for microscopy, genomic sequencing, and bioassays.

### 2. Light microscopy

An aqueous suspension of polyhedral occlusion bodies was examined under a light microscope (Axioscope 40, Carl Zeiss, Germany) with oil immersion at 1000 × magnification. Nomarski contrast was employed for better visualization of the shape of the occlusion bodies.

### 3. Electron microscopy

The extracted occlusion bodies were fixed in suspension by the addition of an equal volume of an 8% paraformaldehyde solution. The mixture was incubated at 4°C for 24 h. Then the polyhedrons were washed with Hank’s solution, additionally fixed with a 1% osmic acid solution, dehydrated in increasing concentrations of ethanol and acetone, and embedded into the Epon–Araldite mixture. Semithin and ultrathin sections were obtained on a Reichert-Jung microtome (Austria). Semithin sections were stained with an azure-II solution, viewed under an AxioImager Z1 light microscope (ZEISS, Germany), and areas of interest were selected for subsequent ultrastructural analyses. Ultrathin sections were counterstained with uranyl acetate and lead citrate. Electron microscopy was carried out using a JEM 1400 electron microscope (Jeol, Japan). Photos were taken with a Veleta digital camera (SIS, Germany). Image analysis was performed in the iTEM software (SIS, Germany).

### 4. Complete genome sequencing

Nucleic acids were extracted from purified viral polyhedrons using the ExtractRNA reagent (CJSC Eurogen, Russia) in accordance with the manufacturer’s instructions. Two microliters of aqueous glycogen (20 mg/ml) was used as a co-precipitator. The precipitate was dissolved in 30 µl of deionized water and utilized as a template in a reverse-transcription reaction to synthesize cDNA according to the SISPA protocol (Sequence-Independent, Single-Primer Amplification) and according to^12^. Synthesized cDNA fragments were purified by means of AMPure beads (Beckman Coulter). The concentration of the DNA was determined with the Qubit dsDNA HS Assay kit on a Qubit 3.0 fluorometer (Thermo Fisher Scientific). Next-generation sequencing libraries were prepared from the samples using the NEB Next Ultra II FS DNA Library Prep Kit for Illumina. Paired-end sequencing (2 × 250 bp) was performed on the MiSeq (Illumina) platform with the MiSeq Reagent Kit v2 (Illumina), resulting in 637–1038-fold genome coverage.

Genome assembly was conducted in the MIRA (v.4.9.6) assembler with default parameters. Multiple-sequence alignment of CPV genomes was performed using the MAFFT (v7.407) algorithm. Maximum likelihood trees were inferred by means of RAxML-NG^13^ within the GTR (G+I) model. Branch support was estimated by the bootstrap method (1000 replications).

Prediction of recombination signals and evaluation of their statistical significance were conducted by the 3SEQ^14^ algorithm.

NCBI GenBank accession numbers for the complete genomes of type 1 CPVs used in the study are provided in Supplementary Table S1.

### 5. A bioassay of DsCPV against the main host species: *D. sibiricus*

#### 5.1. Half-lethal dose and median lethal time

We used the population of *D. sibiricus* collected in fir-cedar forests located at the foothills of the Eastern Sayan. For the bioassay, we obtained the next generation (i.e., F1) of the insects under lab conditions. The larvae were reared at 24°C and 40–60% relative humidity under the 16:8 light:dark regime and were fed 2-year-old shoots of the fir *Abies sibirica* Ledeb.

For infection, we conducted a serial dilution experiment. In particular, the following virus concentrations in suspensions were employed: 10^7^, 10^6^, 10^5^, 10^4^, and 10^3^ occlusion bodies per milliliter. To inoculate the larvae by the virus, the droplet feeding method was applied^15^. Second-instar larvae were individually fed with 0.5 µl of a suspension of the virus in a 10% aqueous sucrose solution. For better visualization, the drinking suspension was colored by a food dye. Larvae of the control group were fed a virus-free solution of sucrose. For each dose, we used 30 individuals. Mortality was recorded daily until pupation. The lethal doses required to kill 50% (LD_50_) of the inoculated larvae were determined by means of the *drc* software package^16^. Differences in LD_50_ were considered statistically significant when fiducial limits did not overlap. Median lethal time (LT_50_) was determined by Kaplan–Meier survival analysis followed by the logrank test with Holm–Sidak adjustment.

#### 5.2. DsCPV productivity

For this assay, we used larvae collected in the larch forests of the Baikal region because the stock from the Sayan *D. sibiricus* population was limited in this study. The larvae were reared under the same conditions as described above (except for the host plant) and were fed the shoots of the larch *Larix sibirica* Ledeb.

For infection, fifth-instar larvae were individually fed 2 µl of a suspension of the virus in a colored 10% aqueous sucrose solution. The virus concentration in the suspension was 2 × 10^7^ polyhedrons/ml. The infected and control groups contained 30 individuals each. The polyhedrons were isolated from dead larvae by crushing a larvae body in a fixed volume of water. Then, the suspension of the homogenate (without filtration to avoid the loss of occlusion bodies) was diluted, and the occlusion bodies were counted on a hemocytometer at 400× magnification. Viral productivity was determined as the number of polyhedrons per larva.

### Determination of the host species range

The experiment involved larvae of nine species of Lepidoptera from six families (except for the main host). Peroral infection was carried out in two ways: drop-feeding and diet-incorporation. Drop-feeding consisted of feeding the insects a drop of a virus suspension (see Table 1 for each species) in a colored 10% sucrose solution. If the larvae refused the sweet droplet, then the infection was implemented by contamination of their diet, i.e., diet-incorporation. In the latter case, the scheme of infection for all species was as follows: 1 ml of the virus suspension was applied with a brush to feed intended for one group of insects. The amount of infected feed was such that a group of insects could eat it before it dried. After the larvae ate the contaminated feed or drank the droplet, they were placed in containers in accordance with the population density optimal for the corresponding species. Virus concentrations in suspensions during the infection of different species were uniform; however, with diet-incorporation, the concentration was higher than that employed for the drop-feeding in order to adjust the regimen for the losses seen in this method when the larvae did not finish their feed.

**Table 1.**
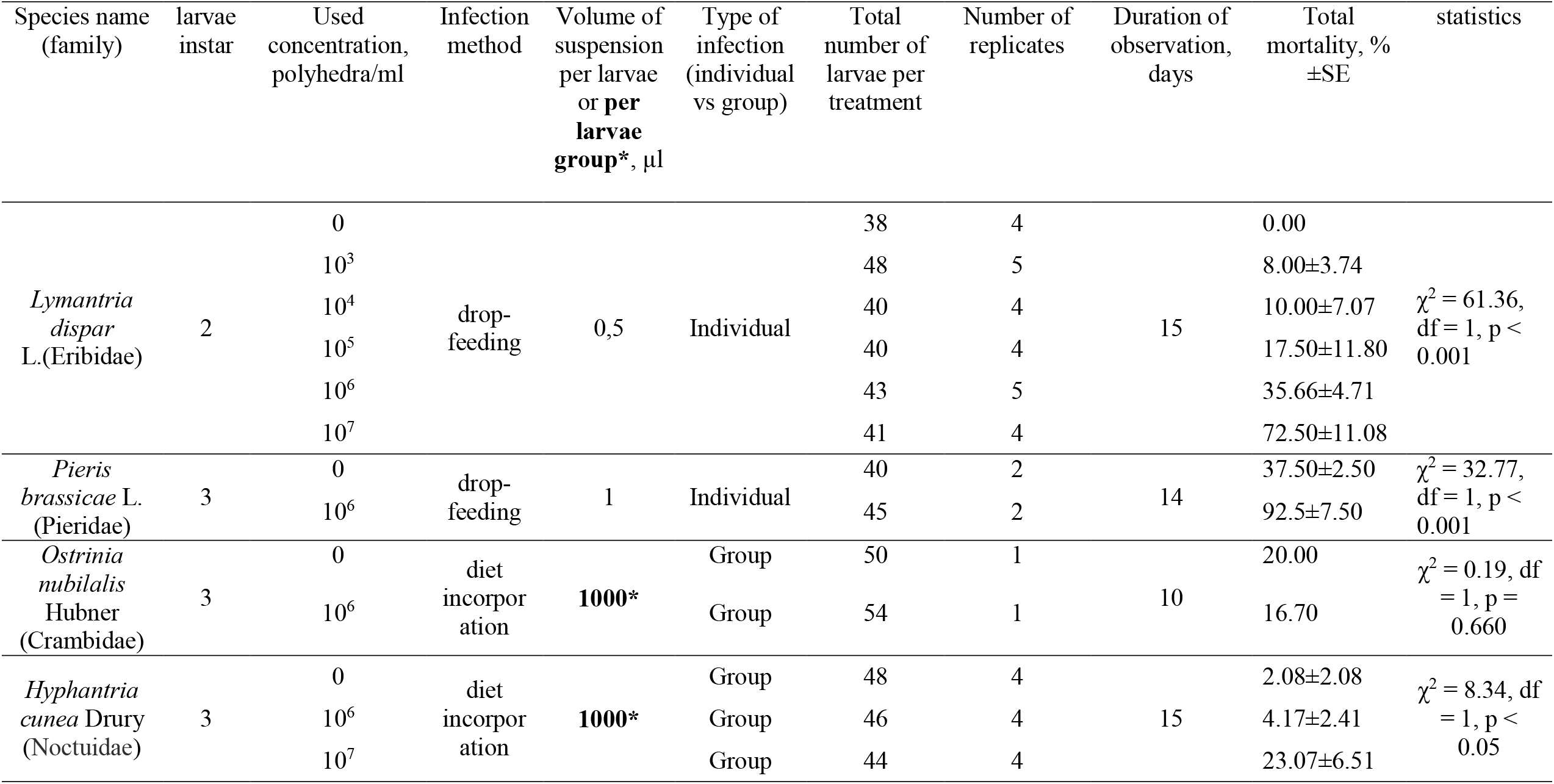
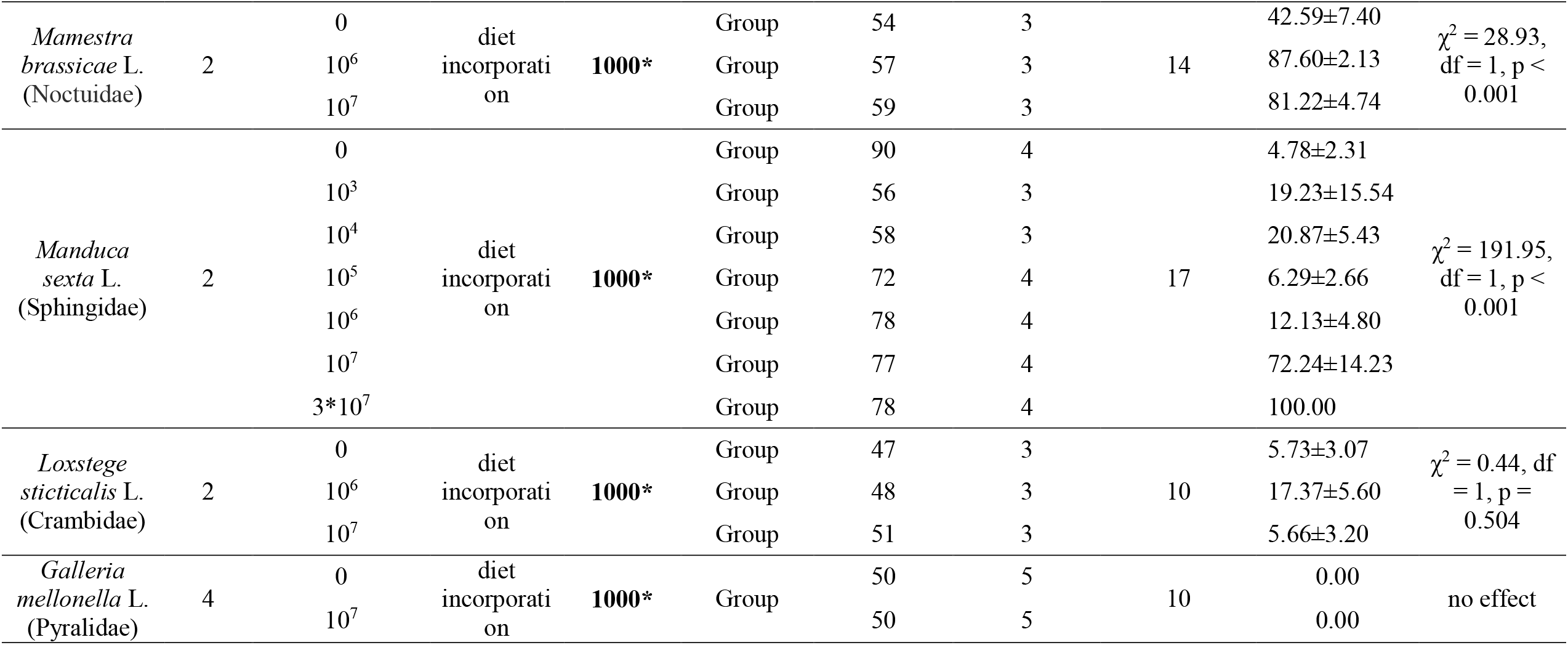
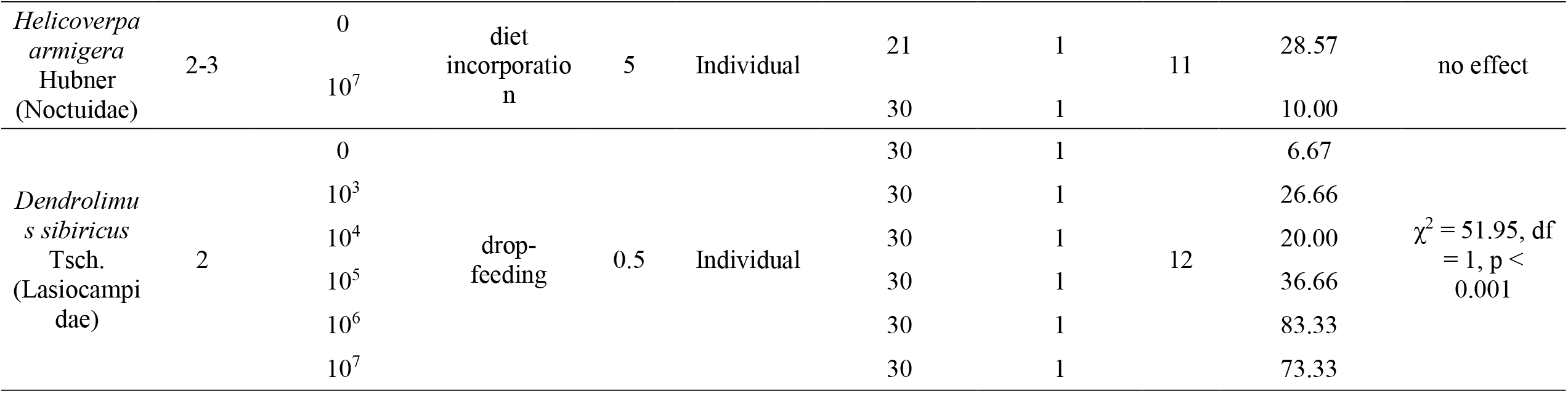
The Bioassay of *Ds*CPV 1 and condition of challenging during host range study

Species, larvae instar, concentration of the virus in a suspension, group assignment of insects, and the number of assay days are shown in Table 1.

#### 6.1. Lymantria dispar

*L. dispar* eggs were collected in the spring from wild birch stands in Novosibirsk Oblast, Russia. Next, the eggs were kept in a refrigerator at 4°C during winter diapause. The hatching of larvae was synchronized with the budburst of birch leaves (first third of May) according to Martemyanov et al. (2015)^17^. After hatching, the insects were reared on cut branches of the silver birch *Betula pendula* Roth. The larvae were reared at 22° C, at natural humidity and under the natural photoperiod.

#### 6.2. Pieris brassicae

Its eggs were collected in a cabbage field (one of untreated fields) on a farm in Novosibirsk Oblast, Russia. The lower leaves of white cabbage (*Brassica oleracea* L.), grown in the laboratory, served as feed for the insects. Larvae were reared at 24–25° C and 50–60% humidity under the natural photoperiod.

#### 6.3. Ostrinia nubilalis

We used insects from a lab population at the All-Russian Institute of Plant Protection (St. Petersburg, Russia). Rearing of larvae was performed on an artificial diet in accordance with the procedure of Frolov et al. (2019)^18^.

#### 6.4. Hyphantria cunea

Larvae of the fall webworm *H. cunea* were collected on mulberry trees in Krasnodar Oblast (Russia) in August 2021 at the egg stage. Larvae were maintained in plastic 2-liter containers and fed with mulberry leaves at 24°C under the natural light/dark regime.

#### 6.5. Mamestra brassicae

In this experiment, larvae of the 1st (F1) generation of the laboratory population were used. The larvae of the parental population were collected in a cabbage field on a farms in Novosibirsk Oblast, Russia. Lower leaves of white cabbage (*B. oleracea* L.), grown in the laboratory, served as feed for the insects. The insects were reared at 24–25°C and 50–60% humidity under the natural photoperiod.

#### 6.6. Manduca sexta

Insect eggs together with artificial feed and the protocol of rearing were kindly provided by T-REX Co. (Russia). The insects were reared at 24–25°C and 50–60% relative humidity under the 16:8 light:dark regime.

#### 6.7. Loxostege sticticalis

Larvae of the 2nd (F2) generation of the laboratory population were used. The larvae of the parental population were collected on a forest edge in Krasnoyarsk Krai (Russia). The larvae were reared at 24°C and a relative humidity of 40–60% under the 16:8 light:dark regime and were fed burdock *Arctium lappa* L. leaves.

#### 6.8. Galleria mellonella

Insect larvae were kindly provided by the insectarium of the Institute of Systematics and Ecology of Animals, the Siberian Branch of the Russian Academy of Sciences (Novosibirsk). Larvae were reared at 28°C and 60% relative humidity on a 12:12 h light:dark cycle and were fed an artificial nutrient medium. The detailed description is given by Dubovskiy et al. (2013)^19^.

#### Helicoverpa armigera

Larvae of the 1st (F1) generation of the laboratory population were used. Pupae of the parental population were collected in Uzbekistan. The larvae were reared on an artificial diet under natural illumination conditions.

### 7. Challenging the main host *D. sibiricus* by the virus passaged through the *M. sexta* host

#### 7.1. Passaging and virus isolation from *M. sexta*

Middle-instar larvae of *M. sexta* we infected by initial virus via diet-incorporation. The feed was supplemented at the concentration of 10^7^ polyhedrons/ml, 1 ml per 20 larvae. Dead larvae were harvested and crushed, and the virus was purified according to the procedure described in section 1.

#### 7.2. Challenging the main host *D. sibiricus* by the virus passaged through *M. sexta*

Second-instar larvae of *D. sibiricus* were employed for this purpose. We used larvae of a filial generation hatched in the laboratory from the parental population that was collected in fir-cedar forests located at the foothills of the Eastern Sayan. The larvae were reared at 24°C and 40–60% relative humidity under the 16:8 light:dark regime and were fed 2-year-old shoots of the fir *A. sibirica*.

For infection, second-instar larvae were individually fed 10^4^ polyhedrons per larva in accordance with the protocol described above. Larvae of the control group consumed a virus-free solution of sucrose. The experiment involved 14 experimental and 14 control individuals. We could not use more larvae for this bioassay because of the limited number of insects. The highest dose was chosen to qualitatively confirm the effectiveness of the virus against the main host. Mortality was recorded daily until the last individual died.

### 8. Increasing the effectiveness of the virus by optical brightener Blankophor (Miles Inc., Pittsburgh, PA, USA)

Larvae of three lepidopteran species were subjected to this experiment: *D. sibiricus, L. dispar*. Drop-feeding was utilized as the infection method To assess the effect of Blankophor on larval mortality from the viral infection, we conducted experiments with the following groups of larvae: 1) a 10% sucrose solution; 2) a solution containing 0.5% of Blankophor and 10% of sucrose; 3) a series of 10-fold dilutions of virus suspensions from 10^3^ to 10^7^ polyhedrons/[ml of the 10% sucrose solution]; and 4) virus suspensions in a solution containing 0.5% of Blankophor and 10% of sucrose. The data are presented together with data on infection of the same species without the optical brightener.

## Results

### Symptoms of infection in the larvae, and the description of DsCPV

The dead larvae of *D. sibiricus* were collected in an outbreak part of a pest’s range in the taiga zone (near the Eastern Sayan mountain; 55.10° N 96.58° E) in 2020. Light microscopic analysis of the dead larvae revealed many polyhedral occlusion bodies with a size of 1.5–2.0 µm (Fig. 1a,b). The dead larvae did not get liquefied, indicating the absence of NPV (Fig. 2a). Transmission electron microscopy confirmed the presence of spherical virus particles (with the average size of 50–60 nm) occluded by a crystal protein that is typical for CPV. The polyhedrons had different shapes including icosahedral (Fig. 1c). The infection of the main host (second-instar larvae of *D. sibiricus*) showed LD_50_ of 68 polyhedrons per larva at 12 days (Fig. 3a). It should be noted that even at the lowest dose of DsCPV (0.5 polyhedrons per larva), we registered a larval mortality rate of 53% throughout the whole larval stage (Fig. S1). Mature polyhedrons were also detectable at a high concentration in the feces of the infected larvae. Median productivity of DsCPV during the infection of middle-instar larvae was 10^8^ polyhedrons/larvae with a range of 2.5 × 10^7^ to 3.1 × 10^8^.

**Figure 1.**
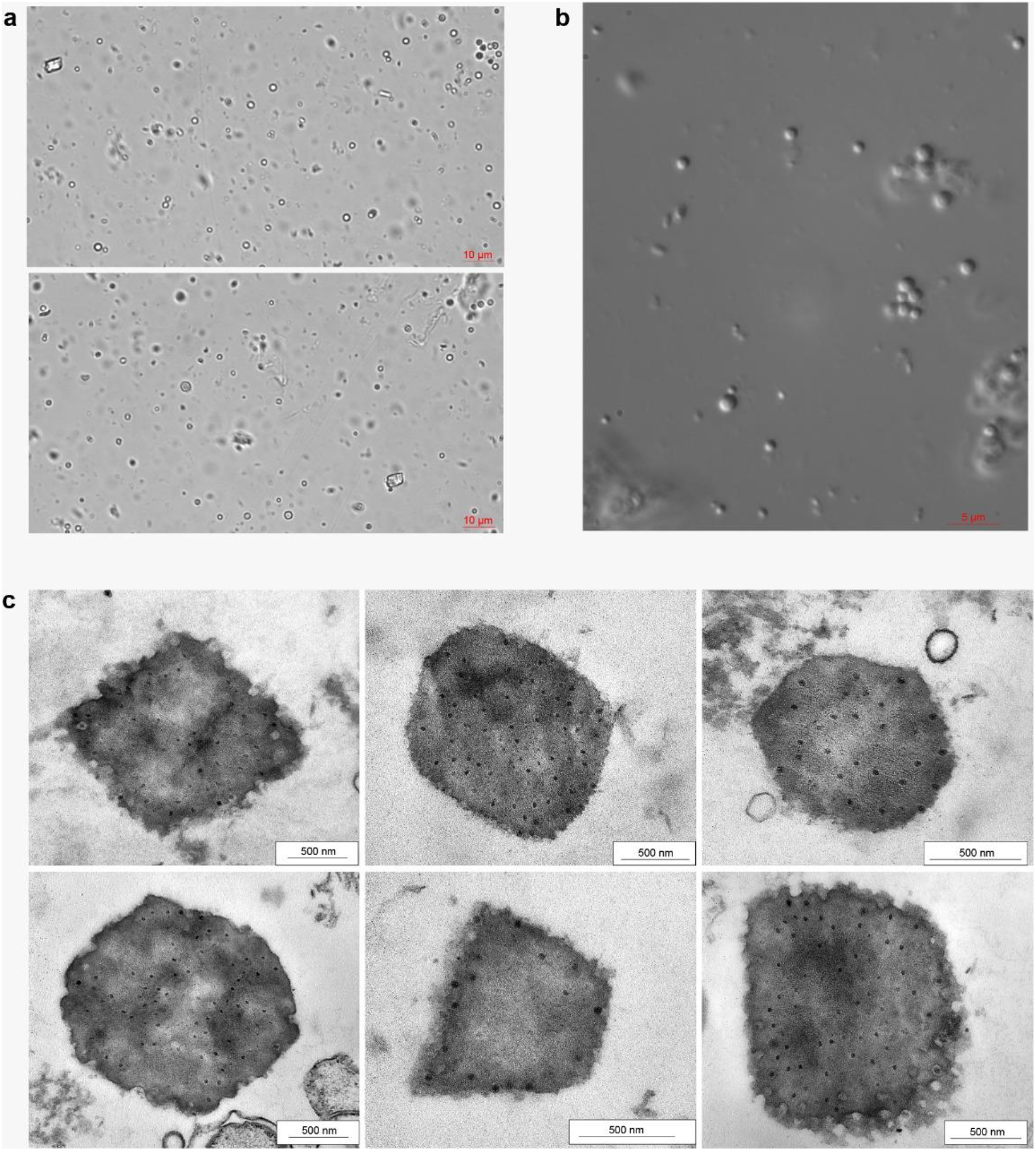
Micrographs of DsCPV1: a) light microscopy (arrows indicate polyhedrons); b) light microscopy with Nomarski contrast; c) transmission electron microscopy of polyhedrons of different shapes (arrows indicate spherical virions typical for the CPV genus).

**Figure 2.**
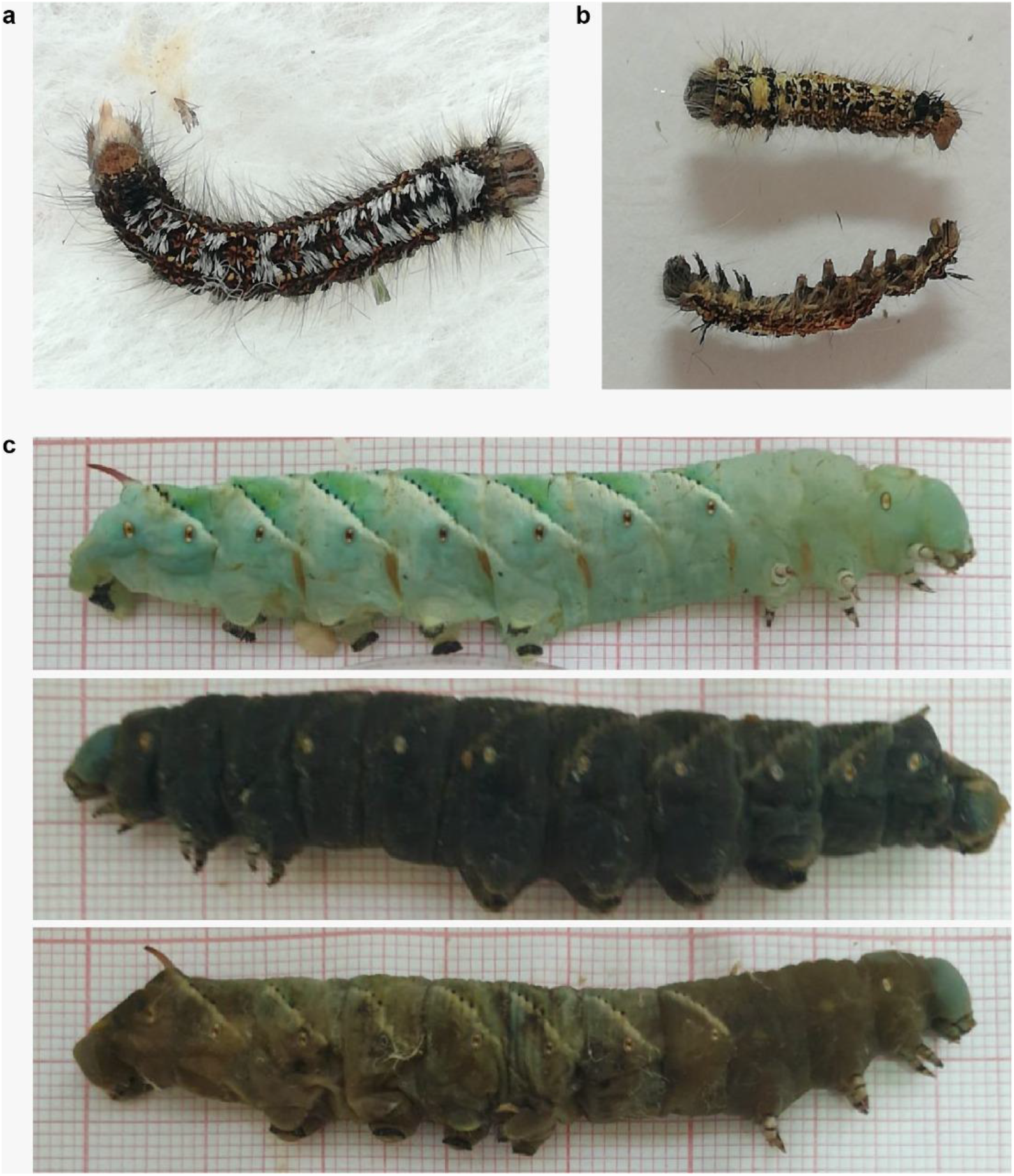
Larvae killed by DsCPV. a) Middle-instar *D. sibiricus* larva; b) young *D. sibiricus* larvae killed by the bioassay; c) dead late-instar *M. sexta* larvae.

**Figure 3.**
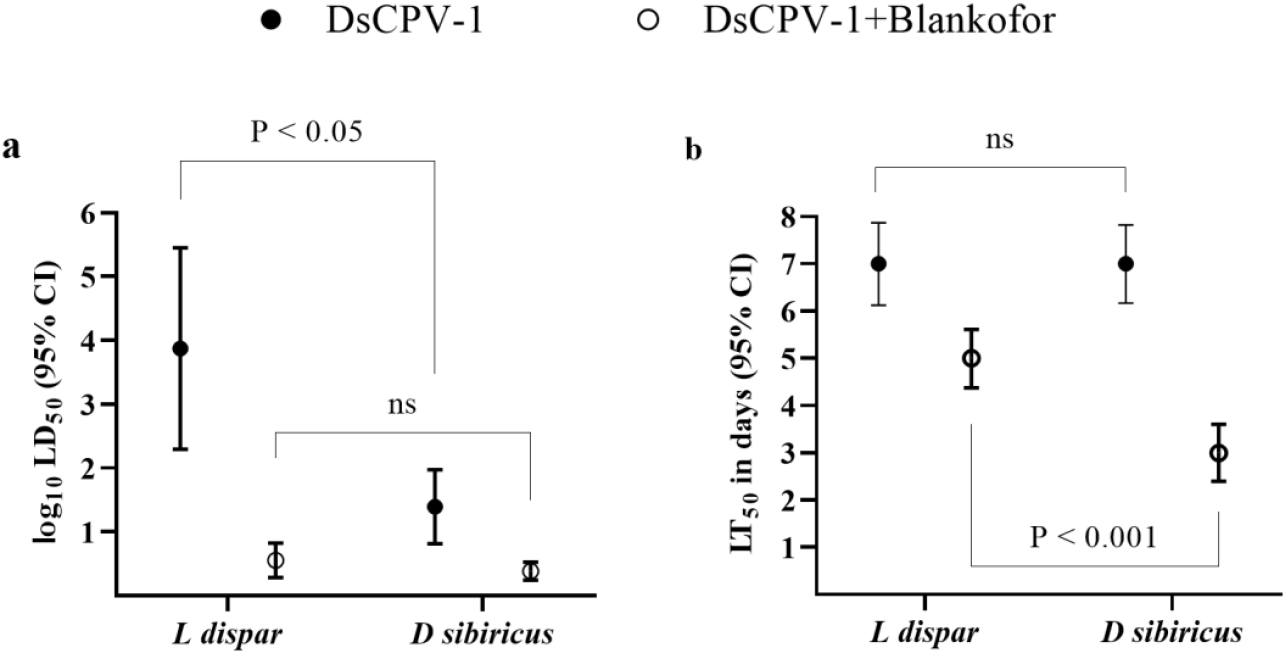
The virulence of DsCPV1 toward second-instar larvae of *D. sibiricus* and *L. dispar*. a) LD_50_ of DsCPV1 combined with the optical brightener (Blankophor); b) LT_50_ of DsCPV1 combined with Blankophor

Just as in other known type 1 CPVs, the sequenced double-stranded-RNA genome of DsCPV1 is composed of 10 linear segments, each containing one ORF (Table S1). According to the phylogenetic analysis of RdRp (Figure 4), which is the only conserved protein domain across RNA viruses and therefore is used to infer their evolutionary relationships^20^, there are only subtle differences between DsCPV1 and both *D. punctatus* CPV 1 (DpCPV1) and one of isolates of *L. dispar* CPV 1 (LdCPV1). Although overall, the genome is highly similar to that of DpCPV1, multiple-sequence alignment of five known CPV1 genomes was rather mosaic: e.g., the third segment of DsCPV1 is much closer to LdCPV1 than to DpCPV1 (Fig. 4). Together with the corresponding incongruence between phylogenies of individual segments (Figures 4 and S1), this finding indicates that some recombination events have occurred (horizontal transmission) between different CPV strains. Accordingly, the 3SEQ analysis of mosaicism predicted that the third segment of DsCPV1 is most likely a result of recombination between DpCPV1 and LdCPV1 (p = 3e-276).

**Figure 4.**
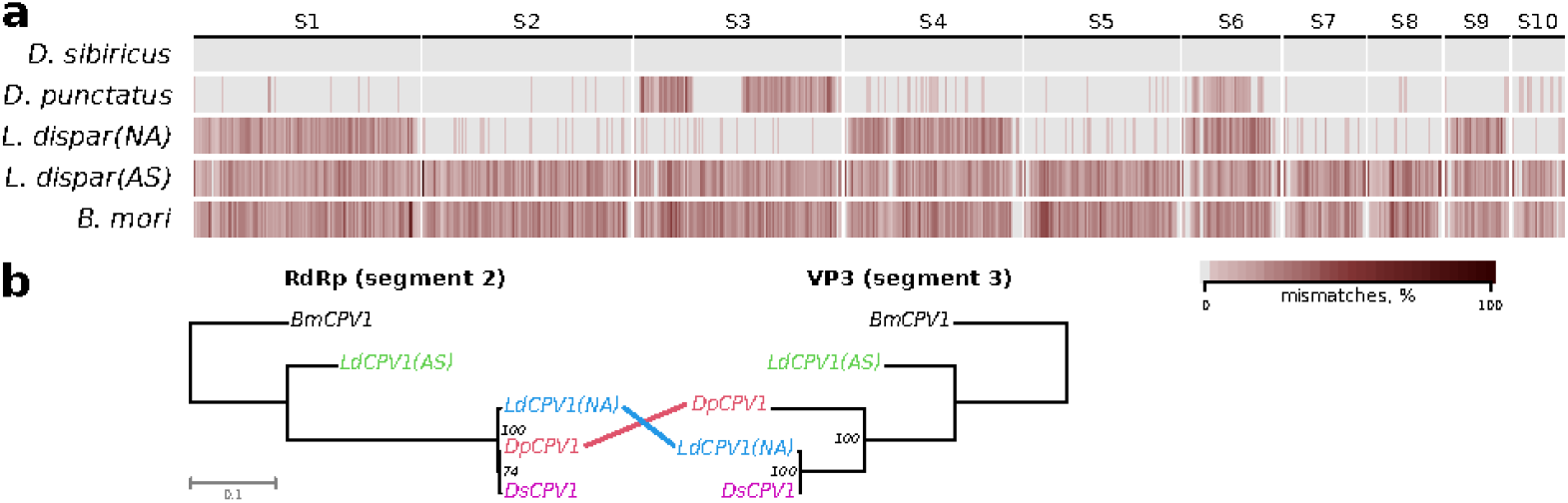
(a) A heatmap of genome-wide sequence differences between *D. sibiricus* CPV1 (DsCPV1) and available complete genomes of other type 1 CPVs. Color intensity corresponds to the proportion of mismatches per genomic window, as defined in the color key. (b) Topological incongruence between the Maximum likelihood estimation trees of segments 2 and 3 encoding RNA-dependent RNA polymerase (RdRp) and structural protein VP3, respectively. The numbers near the branches are Felsenstein’s bootstrap support.

### Examination of the CPV host range and the effect of the optical brightener on CPV potency

The qualitative bioassay of other potential lepidopteran hosts revealed that *P. brassicae, M. brassicae, H. cunea, L. dispar*, and *M. sexta* are susceptible to DsCPV1 infection, whereas *O. nubilalis, G. mellonella, L. sticticalis*, and *H. armigera* are not affected by the virus challenge (Table 1). After DsCPV1 was passaged through *M. sexta* larvae (as the most attractive host for mass production of the virus), reisolated, and fed to *D. sibiricus* larvae, this virus preserved its pathogenicity toward the main host: infection by 10^4^ polyhedrons per second-instar *D. sibiricus* larva killed all the tested larvae. We also compared this virulence of DsCPV1 with its virulence toward another alternative host: *L. dispar*. We demonstrated that the susceptibility of *L. dispar* to DsCPV1 was significantly lower than that of the main host (Fig. 3a,b).

DsCPV1 manifested high effectiveness with the optical brightener. When the adjuvant was added to a DsCPV1 suspension at a concentration of 0.5%, it reduced LD_50_ values for both tested host species, although for *L. dispar* this reduction was much more pronounced, and LD_50_ for mixtures with adjuvants were eventually comparable between the two host species (Fig. 3a). The addition of the brightener also positively affected the speed of killing of the tested hosts. This effect was more significant for the main host (the speed was reduced more than twofold) as compared with the alternative host (Fig. 3b).

## Discussion

Our results clearly indicate that the genus CPV contains some species that hold extra promise for pest management owing to their high virulence to the main host, high productivity (data of this study) even though this virus replicates only in midgut tissue^5^, and broad specificity to lepidopteran host species. The assessment of cumulative mortality of *D. sibiricus* larvae (main host) revealed that even at a dose as low as 0.5 polyhedrons per larva (i.e., only half of the individuals received a viral inoculum), the virus can kill half of the treated population. This high level of infectivity could be a sequence of high sensitivity of the main host or/and successful horizontal transmission of the virus between individuals owing to the presence of mature occlusion bodies in host feces, as we observed in this study and as reported earlier^21^. The virulence of the new strain, DsCPV1, is much higher than that of another closely related strain, CPV1 (derived from other hosts^7, 22^), and than the virulence of the CPV genus in general^22, 23, 24, 25, 26, 27^. One of the most important findings in this study is the unique feature of DsCPV1: an extra wide host range within the order. DsCPV1 successfully infected six of 10 assayed lepidopteran species belonging to five families (Erebidae, Sphingidae, Pieridae, Noctuidae, and Lasiocampidae) by initiating normal pathogenesis in a susceptible host. The virulence of DsCPV1 toward alternative hosts was not as high as that toward the main host but is still comparable with the virulence of other entomopathogens used in pest management^10^. It is known that representatives of the CPV1 taxon have been isolated from different lepidopteran hosts, such as *D. punctatus*^28^, *Bombyx mori*^29^, and *L. dispar*^7^, but until our study, there has been no confirmation of such a wide host range for CPV1 or even for the Baculoviridae family. Phylogenetic analysis of the sequenced DsCPV1 genome showed that it most likely has arisen as a consequence of a recombination event between strains DpCPV1 and LdCPV1, involving a replacement of the third segment of the DpCPV1 genome with its LdCPV1 homolog. It was recently reported that such evolutionary events as reassortment and intragenic recombination are widespread among RNA viruses, including CPV, and serve as a mechanism of adaptation to changing environmental conditions, including host range expansion^30^. The possibility of such an evolutionary mechanism is further supported by evidence that coinfection with several strains of CPV occurs in natural populations^7, 31^. It seems plausible that a recombinant strain—derived from strains that infect different hosts—has acquired a selective advantage in terms of broadening of the host range.

This fundamental finding may form the basis for the following practical implications. First, the found strain DsCPV1 may be considered a quasi-universal agent for population management of economically important lepidopterans pests. This means that a universal scheme of production of a viral insecticide may be implemented to create a viral product for a broad pest control market. This will positively affect the price of the insecticide and its profit margin. Second, it is possible to organize mass production of this strain in vivo in most economically suitable cheap host species (the current problem with mass cultivation of entomopathogenic viruses in cell lines [the cost of the culture medium] is not solved yet). For example, in our study, *M. sexta* proved to be a better candidate for this purpose because it has large quickly developing larvae, does not have diapause, and can be reared on the relatively cheap standard diet. The passaging of DsCPV1 via *M. sexta* (i.e., an alternative host) preserved the pathogenicity toward the main host. We did not quantitatively analyze the virulence against *D. sibiricus* after the passaging via *M. sexta* in this study because of the limited number of *D. sibiricus* larvae available for the study. Even though the passaging through an alternative host will decrease the virulence toward the main host or another susceptible lepidopteran species, the virulence will still be acceptable as long as the cost of virus propagation is attractive for its production. Thus, all the above findings about DsCPV1 should considerably increase the flexibility of viral-insecticide production and are suggestive of good prospects of DsCPV1 as a quasi-universal biological insecticide against lepidopteran pests because this virus has a true regulatory effect on pest population dynamics.

Our data are consistent with the results of another study^11^, which indicates that adding optical brighteners to CPVs significantly (by several orders of magnitude) enhances a virus’s virulence toward the host and expands the host range^27, 32^. Moreover, the applied brightener may increase the susceptibility of an alternative host (which normally has low sensitivity) up to the average susceptibility of the main host (which is normally very high). Thus, the use of an optical brightener(s) at low concentrations as an adjuvant for DsCPV1 should widen the range of target hosts with reasonable virulence toward them. Moreover, the optical brightener here significantly decreased LT_50_ of the hosts, and this parameter is also important for the protective ability of pest control products. As mentioned in the Introduction, at present, there are microbial products on the pest control market that possess a similar set of features. These are dominant insecticides based on *B. thuringiensis*^2^. According to general knowledge about CPV characteristics, i.e., good capacity for vertical transmission^33^, fairly good resistance to environmental factors owing to the polyhedral crystal protein^34^, relatively fast infectious-disease course, and a quick antifeedant effect on the host owing to gut damage^5, 34^, our new highly virulent strain DsCPV1 may be a viable competitor to *B. thuringiensis* products and could be more attractive than narrow-specificity Baculoviridae products. Moreover, owing to vertical transmission via surviving hosts, this strain may cause transgenerational downregulation of pest population size (which is impossible for Bt-based products). This ability is crucial for the control of polyvoltine insect pest species.

In conclusion, we would like to draw virologists’ attention to CPVs in general and to the CPV1 groupe in particular for research on the host range of these viruses (of course, our work was limited to local lepidopteran species), their safety for nontarget species (although all hosts of CPVs are still within the insect class^5^), and molecular mechanisms of host–pathogen interaction. In this report, we provide substantial proof that some representatives of CPV are promising candidates for practical applications, which will help to create an environmentally friendly tool for pest management at a reasonable price in the future.

## Acknowledgment

This study was supported by the Russian Science Foundation (grant # 20-64-46011 to V.V.M. for studying the morphology of the virus and its interaction with *H. cunea*, grant # 21-46-07005 to V.V.M. for investigating the interaction of the virus with *L. dispar, M. sexta, G mellonella, H. armigera*, and *M. brassicae*, and for testing the effect of optical brighteners. Grant of the Ministry of Higher Education and Research of Russian Federation (# FZZE-2020-0026) was used for bioassay of another host species.

## Figure captions

**Figure S1.**
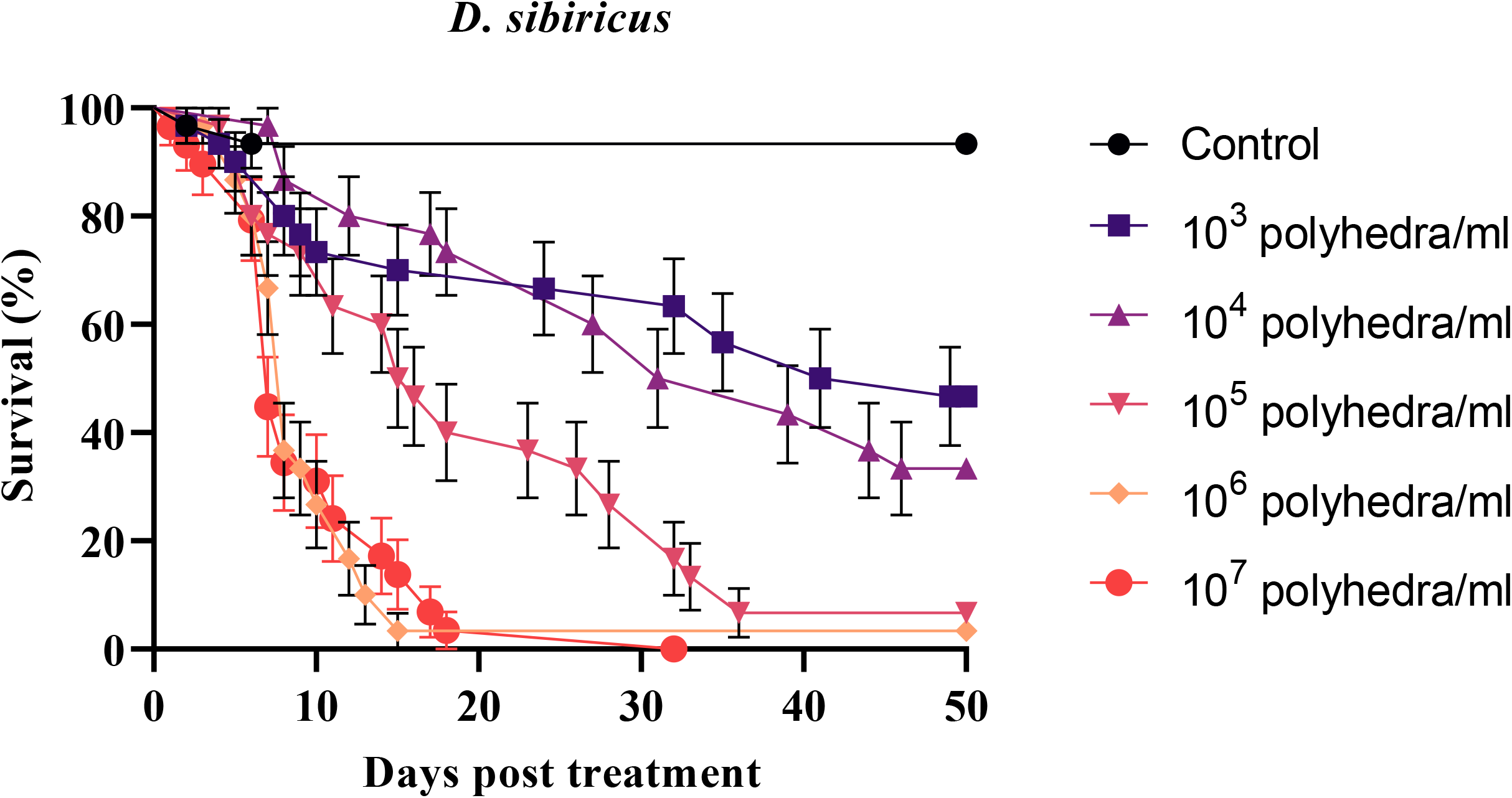
Mortality dynamics of *D. sibiricus* larvae after infection with DsCPV1

**Table S1.**
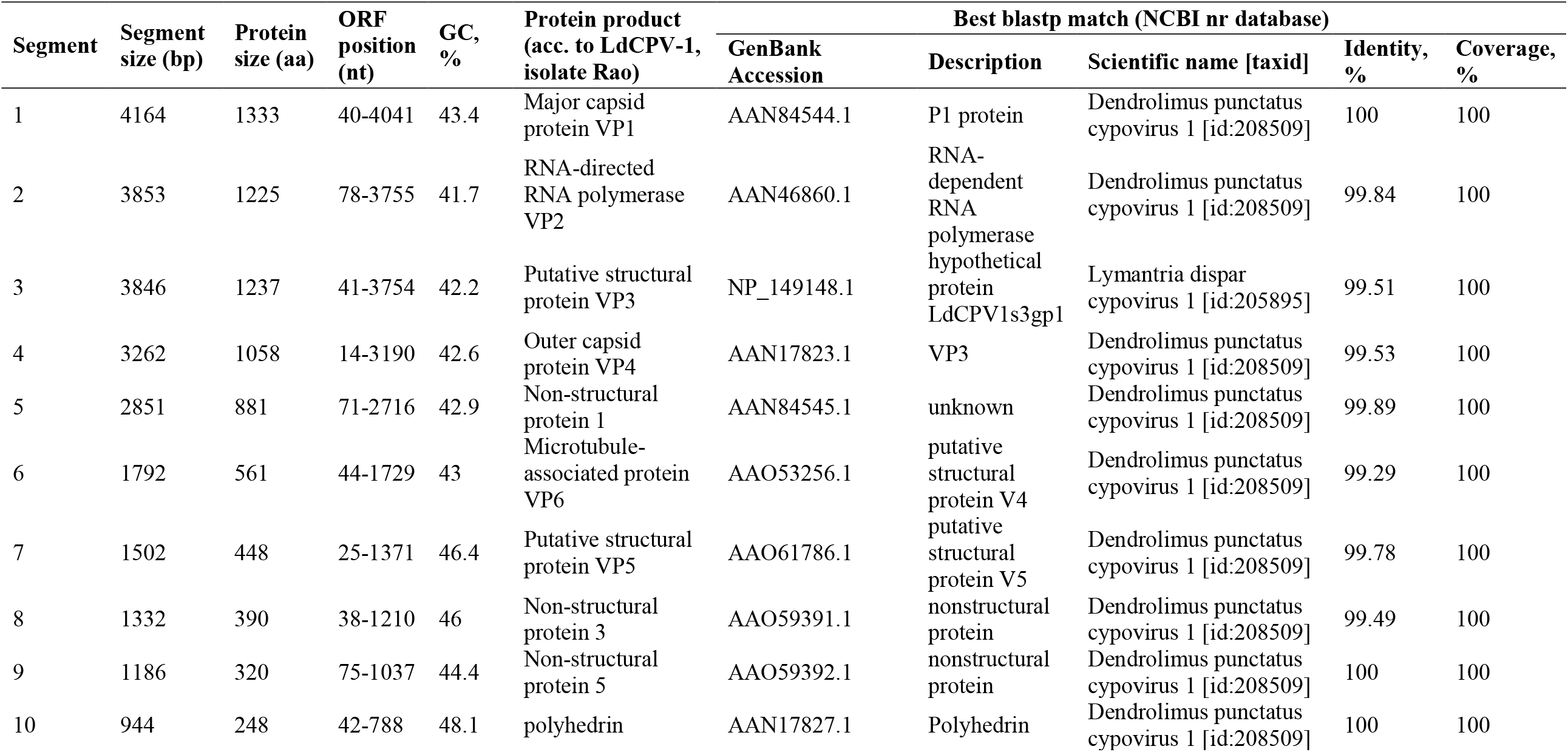
Summary statistics for the assembled DsCPV1 segmented genome.

